# A transcript-wide association study in physical activity intervention implicates molecular pathways in chronic disease

**DOI:** 10.1101/260398

**Authors:** Kajal Claypool, Chirag J Patel

## Abstract

**Background:** Physical activity is associated with decreased risk for several chronic and acute conditions including obesity, diabetes, cardiovascular disease, mental health and aging. However, the biological mechanisms associated with this decreased risk are elusive. One way to ascertain biological changes influenced by physical activity is by monitoring changes in how genes are expressed. In this investigation, we conducted a *transcriptome-wide association study* of physical activity, meta-analyzing 20 independent studies to increase power for discovery of genes expressed before and after physical activity. Further, we hypothesize that genes identified in physical activity are expressed in obesity, inflammation, major depressive disorder and healthy aging.

**Results:** Our analysis identified thirty (30) transcripts induced by physical activity (PA signature), at an FDR < 0.05. Twenty (20) of these transcripts, including *COL4A3, CAMKD1, SLC4A5, EPS15L1, RBM33*, and *CACNG1*, are up-regulated and ten (10) transcripts including *CRY1, ZNF346, SDF4, ANXA1* and *YWHAZ* are down-regulated. We find that several of these physical activity transcripts are associated and biologically concordant in direction with body mass index, white blood cell count, and healthy aging.

**Conclusions:** powerful approach, we found thirty genes that were putatively influenced by physical activity, eight of which are inversely associated with body mass index, thirteen inversely associated with white blood cell count, and three associated and concordant with healthy aging. One gene was significant and concordant with major depressive disorder. These results highlight the potential molecular basis for the protective benefit of physical activity for a broad set of chronic conditions.

## Background

Physical activity is associated with decreased risk for obesity^1,2^, aging^3-6^, all-cause mortality^7^, cardiovascular disease^8-10^, hypertension^11-13^, age-related diseases^14-19^, type 2 diabetes mellitus^20-23^, esophageal cancer^24^, colon cancer^25, 26^, breast cancer^27^, and anxiety and depression^28^. However, while there is strong evidence of the effectiveness of regular physical activity in the prevention of these chronic and acute conditions and premature death^12^, the molecular mechanisms that underlie this protective effect of physical activity are not fully understood. As chronic conditions such as diabetes and obesity reach epidemic status^29-31^, there is a pressing need to gain better understanding of the molecular mechanisms that govern physical activity. Such an understanding can lead to more effective treatment plans, reducing patient risk for multiple chronic conditions.

In the past decade, researchers have used gene expression microarrays to identify functional changes associated with disease and behavior. In fact, several^32-39^ have used microarrays to measure the differential change in gene expression before and after exercise. However, while these investigations have identified putative gene signatures of physical activity, the results are not consistent across the different studies, potentially attributed to small sample sizes, diversity of tissues, exercise intervention protocol, and variations in demographics of the participants. To address these challenges, we conducted a meta-analysis of existing interventional investigations that assay gene expression before and after physical activity. Our large-scale meta-analysis attempts to identify cross-tissue, cross-exercise-type gene expression changes induced by physical activity. We claim that both anaerobic and aerobic activity interventions induce a common set of gene expression changes across tissues, such as peripheral blood mononuclear cells (PBMCs) and muscle. Second, we claim that the molecular changes induced by physical activity are associated with chronic disease. In total, we meta-analyzed 20 independent studies (n=820). We then queried the role of these genes and their association with obesity, inflammation, major depressive disorder and aging to enhance explanation of the observed therapeutic effects of physical activity.

We identified thirty (30) genes to be significantly differentiated (false discovery rate (FDR) < 0.05) after physical activity intervention, with twenty (20) genes up-regulated and ten (10) genes down-regulated. A subset of these genes were significantly associated and biologically relevant for body mass index, white blood cell count and healthy aging. We found one gene associated and concordant with major depressive disorder.

Our work represents, to the best of our knowledge, the largest meta-analysis identifying changes in gene expression associated with physical activity. This physical activity gene expression signature may offer insights into the pathophysiological mechanisms that underlie the potential therapeutic benefits of physical activity observed for a broad swath of chronic conditions.

## Methods

Our overall method is described in Figure 1 (a). We first identified the transcripts differentially expressed after a physical activity intervention. For this, we conducted a large scale meta-analysis^40-43^ over 820 samples drawn from twenty distinct studies. We next attempted to identify the biological significance of genes associated with physical activity in traits such as body mass index (a risk factor for chronic disease), white blood cell count (a marker for inflammation and risk or result of chronic disease), major depressive disorder, and biological aging.

**Figure 1.**
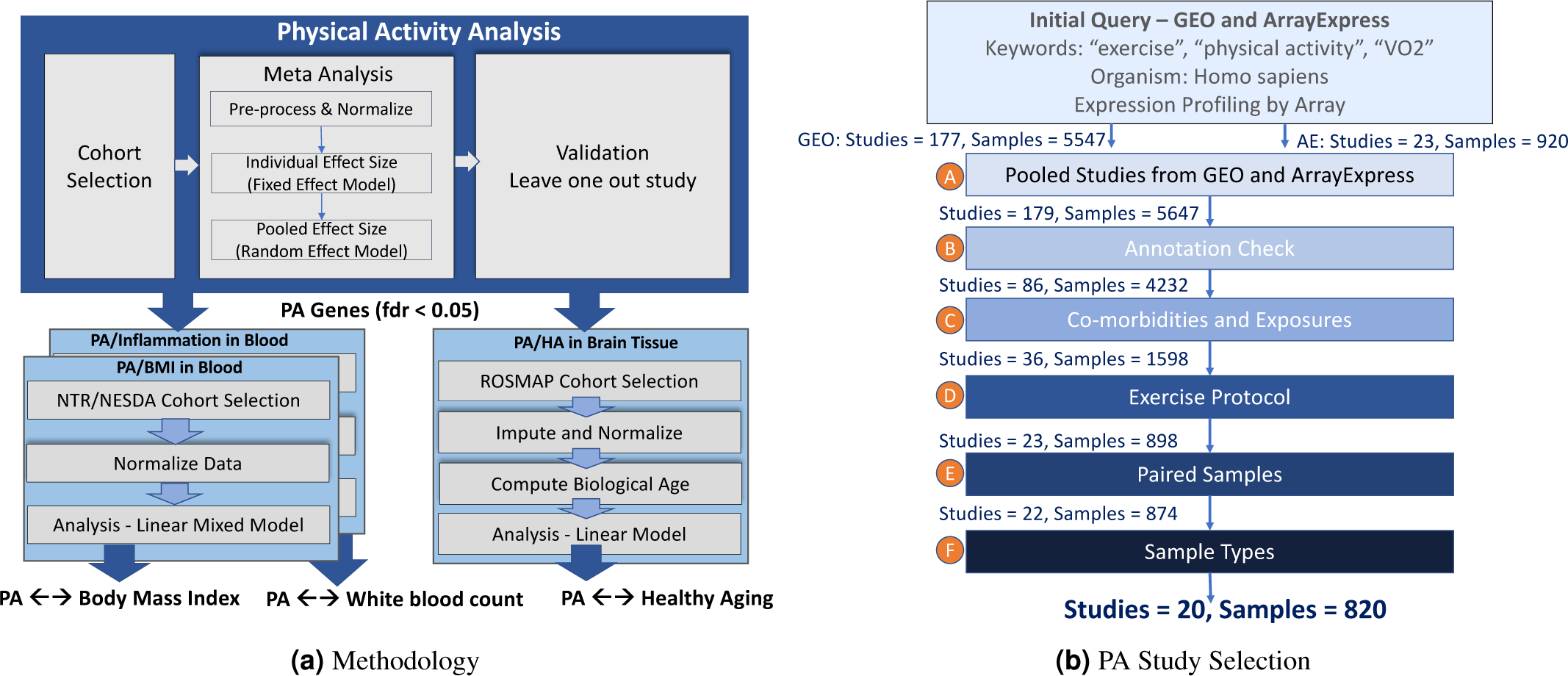
(a) Overall Methodology for the Study, (b) Physical Activity Study Selection - Microarray studies from Gene Expression Omnibus (GEO) and Array Express.

### Data Collection

We cataloged and downloaded gene expression and sample data from twenty (20) publicly available microarray studies that assayed gene expression before and after physical activity in healthy participants. We selected these studies from Gene Expression Omnibus (GEO)^44^ and ArrayExpress^45^ using the query terms `exercise [ALL], physical activity [ALL], VO2 [ALL]` and `Homo sapiens [ORGN]` (to select human gene expression data) and `Expression profiling by array [Filter]` (to select microarray studies). Our search query identified 177 studies in GEO and 23 studies in ArrayExpress. Twenty-one (21) of the ArrayExpress studies overlapped with the GEO studies resulting in a combined set of 179 studies with 5647 samples (Step (A) in Figure 1 (b)). See Supplementary File B for the full list of the 179 studies. We used the rentrez R package^46^ to execute the query on GEO and the ArrayExpress R package^47^ to execute the query on ArrayExpress.

We curated these 179 studies using the selection criteria highlighted in Figure 1 (b). First, to ensure uniform transcript-level analysis, we verified that there were known probe to transcript annotations based on GPL annotation and the information available in GEOmetaDB^48^. This resulted in a total of 86 studies with 4292 samples (Step (B) in Figure 1 (b)). Next, we eliminated studies that included co-morbidities and/or exposures prior to exercise to minimize confounding. Fifty studies with a total 2694 samples were filtered out in this step (Step (C)). In Step (D), we verified that the investigation either intervened with aerobic (AR) exercise intervention, defined as participants achieving 70% to 90% of their peak heart rate and exercised for at least 30 minutes^49,50^; or anaerobic (AN), where participants trained 2 to 3 times per week^51^. We filtered out studies that did not conform to these criteria, leaving 23 studies with a total of 898 samples. Next, to eliminate time-invariant confounding, we selected studies with paired data, that is gene expression before and after physical activity. This resulted in 22 studies and 874 samples (Step (E)). Finally, we selected studies where the sample types corresponded to tissues in GTex^52^. Two studies conducted on natural killer (NK) cells were removed at this last step (Step (F)). Overall, we identified twenty (20) microarray datasets that conformed to our selection criteria with a total of 820 samples (410 paired samples). Table 1 provides overall statistics for the selected studies while Table 2 details the selected studies.

**Table 1.** Meta Analysis study statistics at a glance.

|  | Number of Studies | Number of Samples | Sex Distribution | GPL |
| --- | --- | --- | --- | --- |
| All | 20 | 820 | ♂:471, ♀:176, ?:173 | 96,570,571,6244,10558,11532 |
| Aerobic | 16 | 613 | ♂:369, ♀:160, ?:84 | 96, 570, 571, 6244, 10558 |
| Anaerobic | 4 | 209 | ♂:102, ♀:16, ?:89 | 570, 6244, 11532 |

**Table 2.**
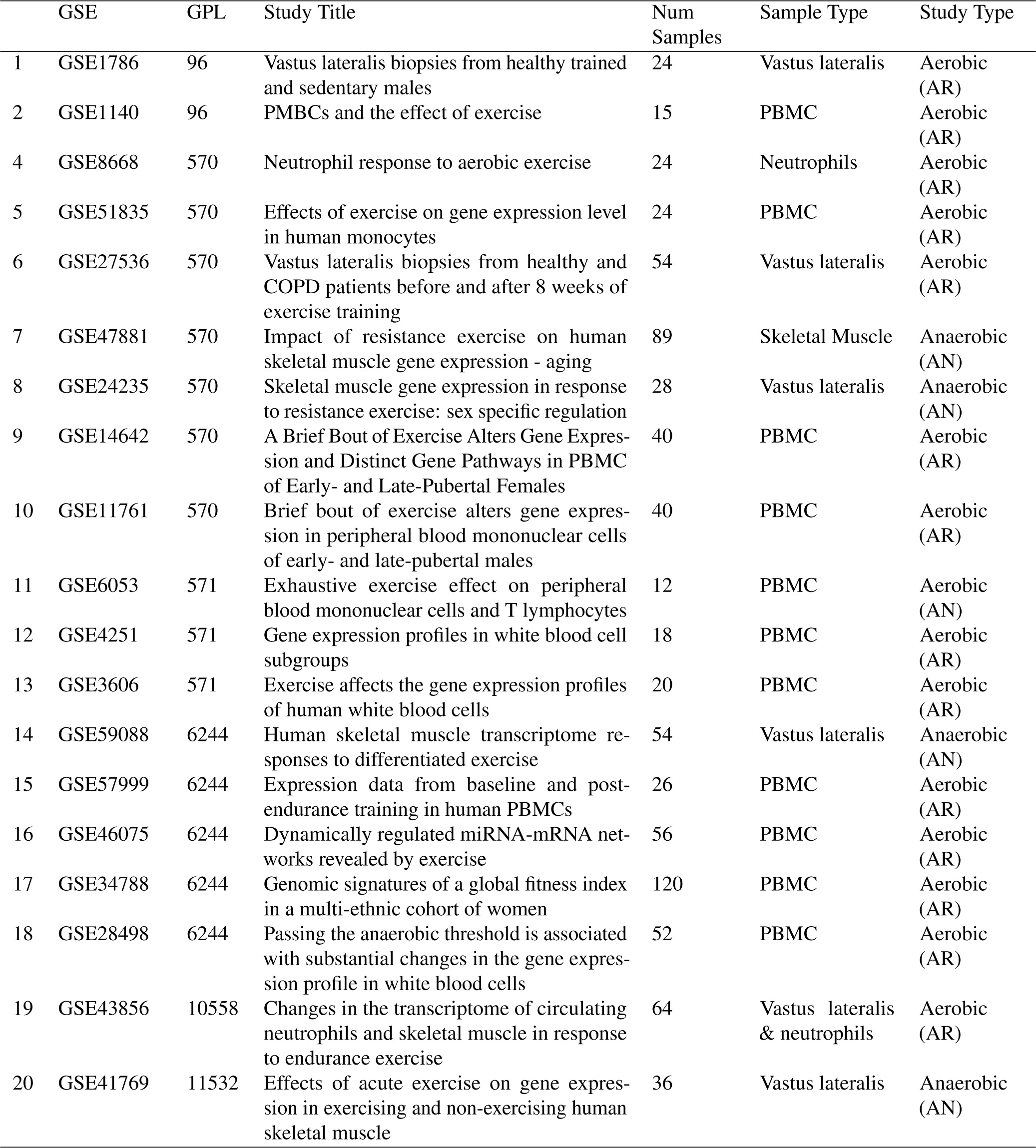
Selected studies from Gene Expression Omnibus. Each study provided paired gene expressions corresponding to before and after physical activity.

|  | GSE | GPL | Study Title | Num Samples | Sample Type | Study Type |
| --- | --- | --- | --- | --- | --- | --- |
| 1 | GSE1786 | 96 | Vastus lateralis biopsies from healthy trained and sedentary males | 24 | Vastus lateralis | Aerobic (AR) |
| 2 | GSE1140 | 96 | PMBCs and the effect of exercise | 15 | PBMC | Aerobic (AR) |
| 4 | GSE8668 | 570 | Neutrophil response to aerobic exercise | 24 | Neutrophils | Aerobic (AR) |
| 5 | GSE51835 | 570 | Effects of exercise on gene expression level in human monocytes | 24 | PBMC | Aerobic (AR) |
| 6 | GSE27536 | 570 | Vastus lateralis biopsies from healthy and COPD patients before and after 8 weeks of exercise training | 54 | Vastus lateralis | Aerobic (AR) |
| 7 | GSE47881 | 570 | Impact of resistance exercise on human skeletal muscle gene expression - aging | 89 | Skeletal Muscle | Anaerobic (AN) |
| 8 | GSE24235 | 570 | Skeletal muscle gene expression in response to resistance exercise: sex specific regulation | 28 | Vastus lateralis | Anaerobic (AN) |
| 9 | GSE14642 | 570 | A Brief Bout of Exercise Alters Gene Expression and Distinct Gene Pathways in PBMC of Early- and Late-Pubertal Females | 40 | PBMC | Aerobic (AR) |
| 10 | GSE11761 | 570 | Brief bout of exercise alters gene expression in peripheral blood mononuclear cells of early- and late-pubertal males | 40 | PBMC | Aerobic (AR) |
| 11 | GSE6053 | 571 | Exhaustive exercise effect on peripheral blood mononuclear cells and T lymphocytes | 12 | PBMC | Aerobic (AN) |
| 12 | GSE4251 | 571 | Gene expression profiles in white blood cell subgroups | 18 | PBMC | Aerobic (AR) |
| 13 | GSE3606 | 571 | Exercise affects the gene expression profiles of human white blood cells | 20 | PBMC | Aerobic (AR) |
| 14 | GSE59088 | 6244 | Human skeletal muscle transcriptome responses to differentiated exercise | 54 | Vastus lateralis | Anaerobic (AN) |
| 15 | GSE57999 | 6244 | Expression data from baseline and post-endurance training in human PBMCs | 26 | PBMC | Aerobic (AR) |
| 16 | GSE46075 | 6244 | Dynamically regulated miRNA-mRNA networks revealed by exercise | 56 | PBMC | Aerobic (AR) |
| 17 | GSE34788 | 6244 | Genomic signatures of a global fitness index in a multi-ethnic cohort of women | 120 | PBMC | Aerobic (AR) |
| 18 | GSE28498 | 6244 | Passing the anaerobic threshold is associated with substantial changes in the gene expression profile in white blood cells | 52 | PBMC | Aerobic (AR) |
| 19 | GSE43856 | 10558 | Changes in the transcriptome of circulating neutrophils and skeletal muscle in response to endurance exercise | 64 | Vastus lateralis & neutrophils | Aerobic (AR) |
| 20 | GSE41769 | 11532 | Effects of acute exercise on gene expression in exercising and non-exercising human skeletal muscle | 36 | Vastus lateralis | Anaerobic (AN) |

We harmonized the gene expression data from each study to include expression data for transcripts that were common to all datasets. All data was normalized and log2 transformed using Robust Multichip Averaging (RMA)^53^ implemented in the Affy package and batch corrected using ComBat^54^.

### Meta-Analysis - Physical Activity Effect Size Computation

Our goal was to compute the pooled/combined effect size for each transcript across the selected twenty studies. For this, we first computed the individual study effect size (corrected Hedges’ g) for each transcript using the paired samples. We used a fixed-effects model to combine gene expression values where multiple probes mapped to the same transcript. We next used a random-effects model to combine effect sizes across the studies. Specifically, we used the DerSimonian-Laird random effects model^55^ wherein the pooled effect size, 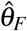, is given as the weighted average of the individual effect effect sizes from each study. That is,

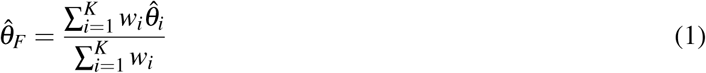

And the weights are given as:

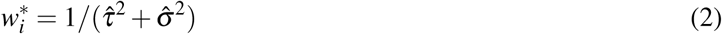

where 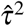 is the measure of the inter-study heterogeneity and 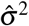 are the variances of each study’s effect size estimate 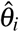 For each pooled effect size, we tested for the null hypothesis of no effect 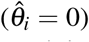, and the null hypothesis of no heterogeneity 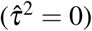. To control for multiple hypothesis testing we computed the Benjamini-Hochberg (BH) false discovery rate (FDR), the estimated proportion of the discoveries made that were false. We used the `R` package `metafor`^56^ to estimate the pooled effect size, the significance and the inter-study heterogeneity. We tested the inter-study heterogeneity using a Chi-square distribution with `k - 1` degrees of freedom.

We next used a two-step method to identify the significant transcripts from the meta-analysis. First, we identified differentially expressed transcripts, 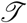. We considered the transcript t ∈ 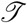 significant if the absolute value of the pooled effect size of the transcript t was greater than 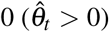, the false discovery rate (FDR) less than 0.05 (FDR_t_ < 0.05) and the inter-study heterogeneity 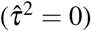 accounted for less than 50% of the variability in point estimates or was not highly significant (p < 0.0001)^57^. To ensure that our findings were not driven by any single study, we next conducted a leave one out sensitivity analysis by iteratively removing one (1) study at a time^42^. Transcripts discovered in Step 1, 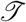, were considered valid and robust, if they had a p < 0.05 across all leave-one-study out analyses. We report physical activity genes surviving the leave-one-out analysis as the PA signature.

### Physical Activity and Phenotype Association

We correlated the PA signature genes with body mass index, white blood cell count and major depressive disorder in monozygotic and dizygotic twins. Finally, we interrogated the association between PA signature genes, measured by RNASeq, and brain tissue epigenetic age, measured by genome-wide methylation arrays. We defined age deceleration (“healthy aging”) as the difference between chronological age and epigenetically predicted age^58^.

### Phenotype Cohorts

We interrogated the Netherlands Twin Registry (NTR) and the Netherlands Study of Depression and Anxiety (NESDA) twin cohort to determine the association between PA signature genes and body mass index, white blood cell count and major depressive disorder. The NTR/NESDA twin cohort provides clinical and gene expression measurements on 2561 twin pairs. Clinical phenotypes available in the data set include demographic variables, age, sex, body mass index (BMI), clinical markers like white blood cell count (WBC) and neutrophil count, and clinical diagnosis of major depressive disorder (MDD), together with information on twin zygosity and family (see Table 3 for summary statistics). The dataset was downloaded from the dbGaP website, under pht002805.v1.p1(https://www.ncbi.nlm.nih.gov/gap/?term=pht002805.v1.p1) and normalized prior to analysis.

**Table 3.** Summary Statistics for the NTR/NESDA and ROSMAP Cohort

| Variable | NTR/NESDA | ROSMAP Cohort |
| --- | --- | --- |
| # Samples | 2561 | 547 |
| Tissue | Blood | Brain |
| Age | 34.69 ± 11.59 | 86.75 ± 5.10 |
| Sex | ♂:881, ♀:1680 | ♂:199 ♀:348 |
| BMI | 23.86 ± 4.28 | - |
| WBC | 6.46 ± 2.06 | - |
| MDD | No MDD:2504, MDD:57 | - |
| Zygosity | MZ:1344, DZ:1217 | Not Applicable |
| Education | - | 16.48 ± 3.47 |
| APOE Mutation | - | Yes:134, No: 413 |

To determine the association between physical activity and brain aging, we used the longitudinal clinical-pathologic study of aging and Alzheimer’s disease (AD) from Rush University (ROSMAP). The ROSMAP dataset is a collection of data from two separate studies conducted by the Rush University. ROS is a longitudinal clinical-pathologic cohort with individuals from 40 different religious orders across the United States. MAP is a longitudinal, epidemiologic clinical-pathologic cohort study of common chronic conditions of aging and Alzheimer’s disease. We merged clinical, RNASeq and DNA Methylation data to analyze the association of PA signature genes with aging. ROSMAP clinical data includes clinical, pathology and demographic data including age at time of death, sex and years of education for 1102 individuals. RNA Seq data, generated from RNA extracted from the gray matter of the dorsolateral prefrontal cortex, contain fragments per kilobase of transcript per million mapped reads (FPKM) for 640 subjects from the ROS and MAP cohorts. RNA Sequencing was done using Illumina HiSeq, and data was normalized and batch-corrected. Lastly, we also made use of the DNA Méthylation data from ROSMAP. DNA méthylation was performed on prefrontal cortex samples collected from 740 individuals using the Illumina HumanMethylation450 BeadChip. Figure 2 outlines the merging steps we used to generate the cohort for age deceleration/ age deceleration analysis. (See Table 3 for summary statistics). ROSMAP study data were provided by the Rush Alzheimer’s Disease Center, Rush University Medical Center, Chicago.

**Figure 2.**
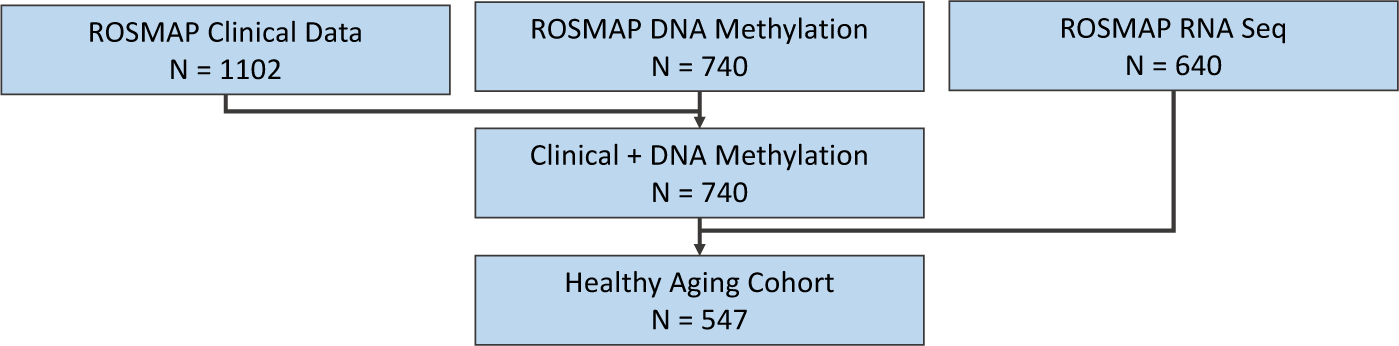
Selection of cohort from ROSMAP for Age Deceleration.

### Associating PA Signature Genes with Phenotypes

#### Linear Mixed Models: Body mass index, White blood cell count and Major depressive disorder

We associated the physical activity transcripts (PA signature genes) in the twin cohort data (NTR/NESDA) with body mass index (BMI), white blood cell count (WBC) and major depressive disorder (MDD). Specifically, we regressed body mass index (BMI) on each of the physical activity genes identified by the leave-one-study out validation step (PA signature genes) (g ∈ *PA Signature*), adjusting for fixed effects of age and sex, and random effects of family and twin monozygosity. One random effect accounted for variation within families and another for monozygotic twins resulting in a linear mixed model^59,60^ of the form:

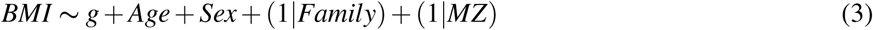

We interrogated the NTR/NESDA twins to identify associations between physical activity genes and white blood cell count, an indicator of inflammation. In addition to age and sex, we adjusted the models for BMI and smoking status to measure the association of physical activity with white blood cell count independent of BMI and smoking status^61-64^. We also adjusted the model for the random effects for family and monozygosity resulting in:

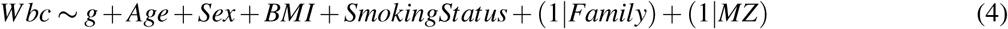

Further, we tested the twin cohort for association between physical activity and major depressive disorder. We adjusted the MDD models for age, sex and alcohol consumption, a known depressant^65^ resulting in models of the form:

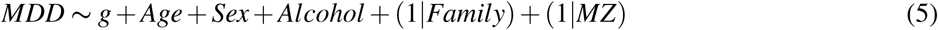

We used the **R** package `lme4`^66^ and `lmerTest`^67^ for our linear mixed model, and `r2glmm`^67^ to compute the R^2^ using the Nakagawa and Schielzeth approach. We report the summary effect size of all PA genes that were found to be significant in our mixed model analysis at p < 0.05.

#### Linear Model: Epigenetic Aging

To perform the association analysis for biological aging, and in particular healthy aging (HA) of the brain, we first computed the epigenetic or biological age for all samples in our healthy aging cohort (N = 547) with 353 epigenetic markers to compute the biological age^58^. Three out of the 353 epigenetic markers were missing (< 1%). We imputed these missing methylation data using knn-imputation^68^ and normalized the DNA methylation data using BMIQ normalization^69^. We verified the predicted epigenetic age with the DNA methylation age calculator at https://labs.genetics.ucla.edu/horvath/dnamage/. We created anew variable, HA, as the difference between chronological age and the estimated epigenetic age. The chronological age for all participants above 90 was hidden in the dataset (set to 90+). We therefore set the age of all individuals above 90 to 91. We then regressed `HA` on each of the PA signature genes using a linear model adjusted for sex and study (ROS or MAP study)^70^ using a linear model of the form:

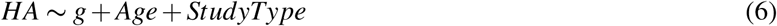

We report the summary effect size of all PA genes that were found to be significant in our analysis at p < 0.05.

For all three phenotypes, we also tested for enrichment of the PA signature genes in the genes associated with the phenotype using a hypergeometric distribution.

For reproducibility, the analysis scripts (code) used for this effort are available in a Git repository (https://kajalc@bitbucket.org/kajalc/physical-activity-analysis.git). Additionally, we have a R Shiny application available to facilitate further exploration of the results at http://apps.chiragjpgroup.org/pagex/.

## Results

### Identifying genes differentially expressed with physical activity

In our transcript-wide analysis of paired gene expression measurements, we found thirty (30) transcripts expressed (FDR < 0.05). Of the 30, twenty (20) were up-regulated and ten (10) transcripts were down-regulated (see Figure 3 and Table 4) after physical activity. Two (2) transcripts, COL4A3 (up-regulated) and SLC25A15 (down-regulated) were found to be significant at FDR < 0.01. All thirty (30) transcripts were found to be consistently associated with physical activity in the leave one study out analysis. Figure 4 shows the individual study and pooled effect sizes of the top four (4) transcripts (by largest absolute effect size). For a comparison of effect sizes of all genes see Supplementary File A.

**Figure 3.**
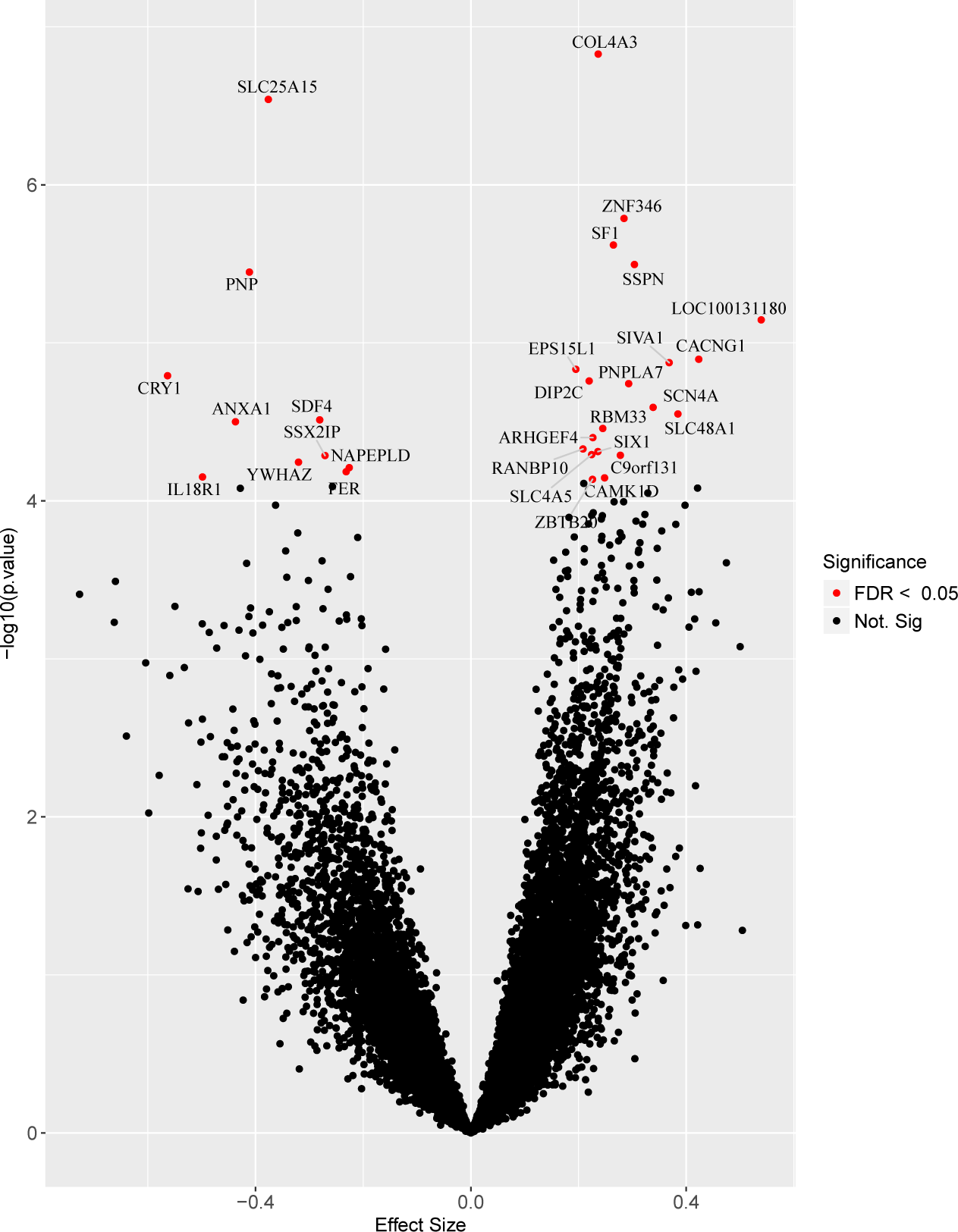
Up- and Down-regulated genes identified by paired meta-analysis of gene expressions taken from blood and muscle tissue (N = 820). Genes significant at false discovery rate < 0.05 are shown in red.

**Figure 4.**
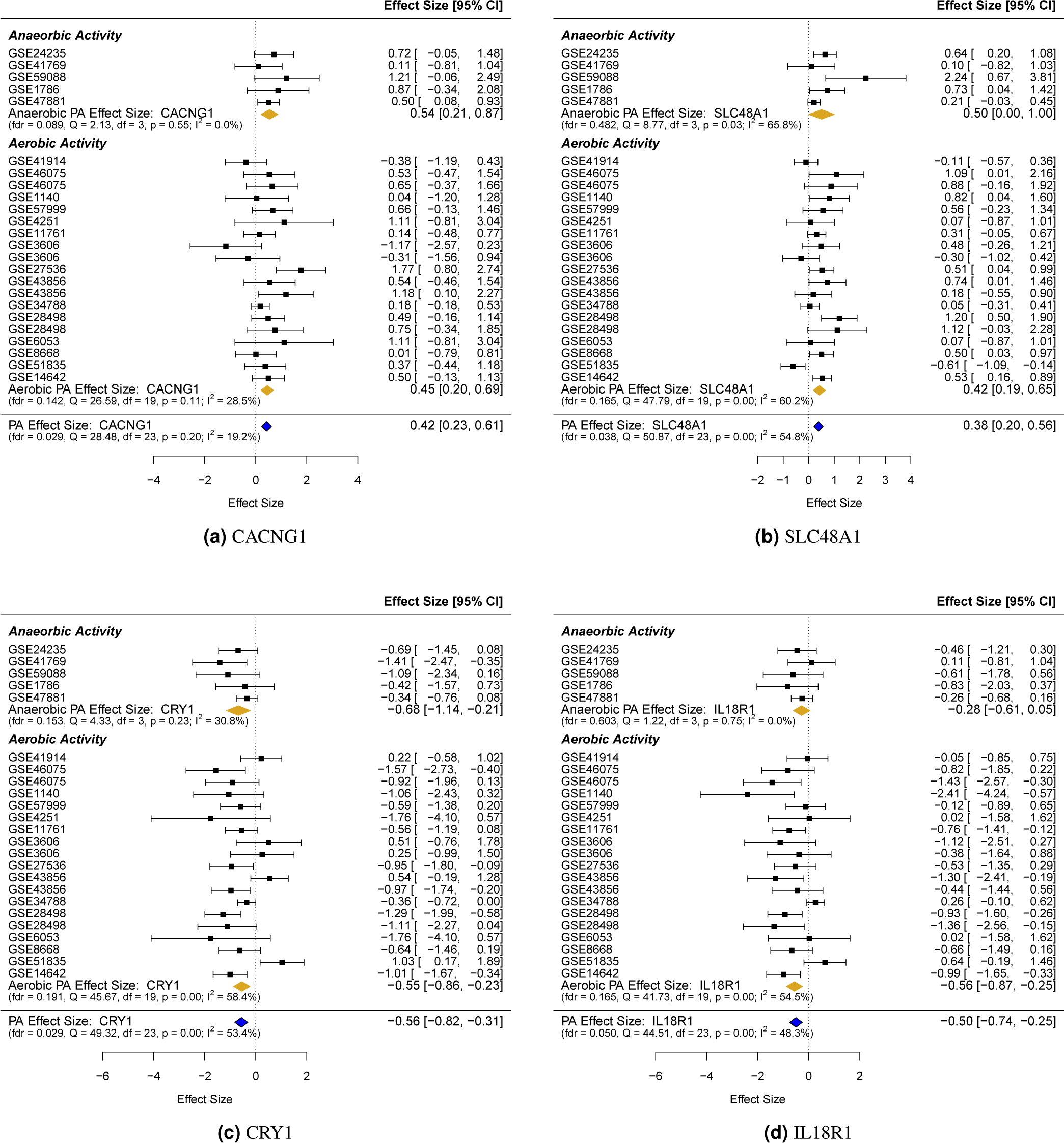
Four differtially expressed genes with biggest effect sizes. For many of the studies, these genes were not significant within the individual studies but were shown to be significant in our meta-analysis.

**Table 4.** Significant PA transcripts. Transcripts differentially expressed as a result of any type of physical activity with FDR < 0.05 and surviving leave-one-out study analysis at p < 0.05.

| Gene | Description | Effect Size | FDR | Direction |
| --- | --- | --- | --- | --- |
| ARHGEF4 | Rho Guanine Nucleotide Exchange Factor 4 | 0.23 | 0.04 | ↑ |
| C9orf131 | Chromosome 9 Open Reading Frame 131 | 0.28 | 0.04 | ↑ |
| CACNG1 | Calcium Voltage-Gated Channel Auxiliary Subunit Gamma 1 | 0.42 | 0.029 | ↑ |
| CAMK1D | Calcium/Calmodulin Dependent Protein Kinase ID | 0.25 | 0.05 | ↑ |
| COL4A3 | Collagen Type IV Alpha 3 Chain | 0.24 | 0.003 | ↑ |
| DIP2C | Disco Interacting Protein 2 Homolog C | 0.22 | 0.03 | ↑ |
| EPS15L1 | Epidermal Growth Factor Receptor Pathway Substrate 15 Like 1 | 0.19 | 0.03 | ↑ |
| LOC100131180 |  | 0.54 | 0.02 | ↑ |
| PNPLA7 | Patatin Like Phospholipase Domain Containing 7 | 0.29 | 0.03 | ↑ |
| RANBP10 | RAN Binding Protein 10 | 0.21 | 0.04 | ↑ |
| RBM33 | RNA Binding Motif Protein 33 | 0.25 | 0.04 | ↑ |
| SCN4A | Sodium Voltage-Gated Channel Alpha Subunit 4 | 0.34 | 0.04 | ↑ |
| SF1 | Splicing Factor 1 | 0.26 | 0.01 | ↑ |
| SIVA1 | SIVA1 Apoptosis Inducing Factor | 0.37 | 0.03 | ↑ |
| SIX1 | SIX Homeobox 1 | 0.24 | 0.04 | ↑ |
| SLC48A1 | Solute Carrier Family 48 Member 1 | 0.38 | 0.04 | ↑ |
| SLC4A5 | Solute Carrier Family 4 Member 5 | 0.22 | 0.04 | ↑ |
| SSPN | Sarcospan | 0.30 | 0.01 | ↑ |
| ZBTB20 | Zinc Finger And BTB Domain Containing 20 | 0.23 | 0.05 | ↑ |
| ZNF346 | Zinc Finger Protein 346 | 0.28 | 0.01 | ↑ |
| ANXA1 | Annexin A1 | -0.44 | 0.04 | ↓ |
| CRY1 | Cryptochrome Circadian Clock 1 | -0.56 | 0.03 | ↓ |
| FER | FER Tyrosine Kinase | -0.23 | 0.05 | ↓ |
| IL18R1 | Interleukin 18 Receptor 1 | -0.49 | 0.05 | ↓ |
| NAPEPLD | N-Acyl Phosphatidylethanolamine Phospholipase D | -0.23 | 0.05 | ↓ |
| PNP | Purine Nucleoside Phosphorylase | -0.41 | 0.01 | ↓ |
| SDF4 | Stromal Cell Derived Factor 4 | -0.28 | 0.04 | ↓ |
| SLC25A15 | Solute Carrier Family 25 Member 15 | -0.38 | 0.003 | ↓ |
| SSX2IP | SSX Family Member 2 Interacting Protein | -0.27 | 0.04 | ↓ |
| YWHAZ | Tyrosine 3-Monooxygenase/Tryptophan 5-Monooxygenase Activation Protein Zeta | -0.32 | 0.04 | ↓ |

### Association with Body Mass Index

Of the thirty (30) identified physical activity genes, seven (7) genes, *C9orf131, CACNG1, EPS15L1, ANXA1, SSX2IP, PNP* and *YWHAZ*, were significantly associated with body mass index (p < 0.05), when adjusted for fixed effects of age and sex, and random effects due to family and monozygosity. We found one additional gene, *SSPN*, to be marginally significant (p = 0.06). Furthermore, all genes were in biologically concordant directions (i.e., down-regulated in PA and up-regulated in BMI and vice versa). Specifically, increased expression of up-regulated PA signature genes, *C9orf131, CACNG1* and *EPS15L1* were all associated with lower BMI. A one unit increase in the expression of *EPS15L1*, for instance, was associated with a -0.35 unit decrease in BMI (95% CI: -0.586, -0.114). Increased expression of down-regulated PA signature genes *ANXA1, PNP, SSX2IP* and *YWHAZ*, on the other hand, were associated with increased BMI. We found that *SSX2IP* is associated with the largest increase in BMI, with a one unit increase in the expression of *SSX2IP* associated with a 0.41-unit increase in BMI (95% CI: 077, 0.743). Figure 5 (a) summarizes the effect sizes of the PA signature genes significantly associated with BMI. All eight PA signature genes were inversely associated with body mass index. The variance explained in body mass index by physical activity genes was 8% (95% CI: 0.066, 0.106). Removing physical activity genes from the model significantly reduced the goodness of fit, as indicated by likelihood ratio test - effect of PA genes: χ^2^ (30) = 67.41, p < 6.78 *e*^−05^. A hypergeometric test to evaluate the enrichment of PA signature in BMI was nominally significant (P (x ≥ 8) = 0.04).

**Figure 5.**
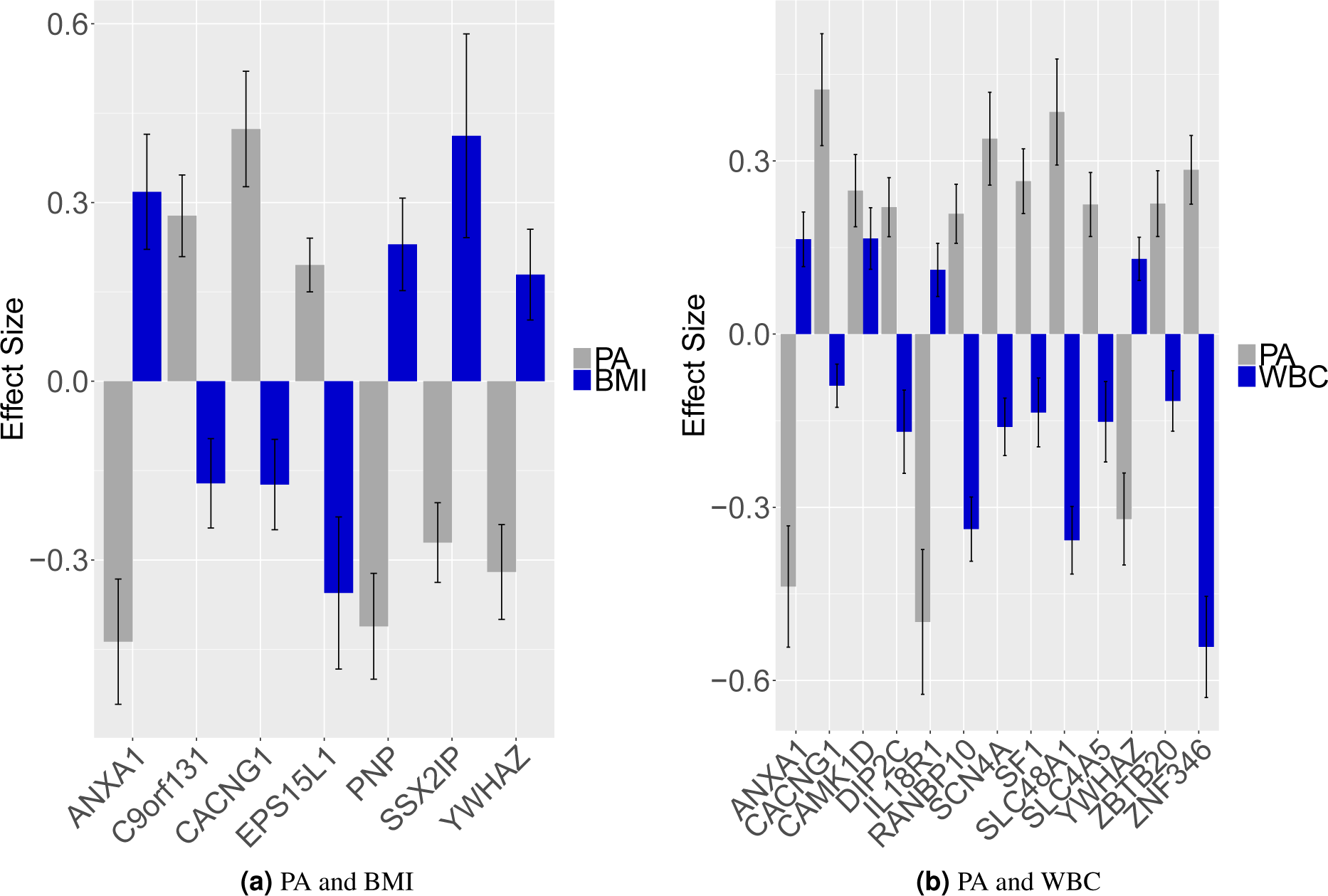
(a) Association of PA signature genes with body mass index. (b) Association of PA signature genes with white blood cell count.

### Association with White Blood Cell Count

We found thirteen (13) PA signature genes to be significantly associated with white blood cell count (p < 0.05), when adjusted for fixed effects of age, sex, BMI, smoking status, and random effects of family and monozygosity. Twelve of these significantly associated transcripts were in biologically concordant directions with physical activity. Specifically, we found increased expressions of up-regulated physical activity genes, *CACNG1, DIP2C, RANBP10, SCN4A, SF1, SLC48A1, SLC4A5, ZBTB20* and *ZNF346*, were associated with a decrease in white blood cell count. A one unit increase in the expression of *ZNF346*, for example, was associated with a 0.54 decrease in white blood cell count. Increased expressions of down-regulated PA signature genes, however, were associated with increased counts of white blood cells, with for example, a one unit increase in *YWHAZ* expression associated with a 0.13 increase in white blood cell count. Figure 5 (b) summarizes the effect sizes of PA signature genes significantly associated with white blood cell counts compared with PA signature gene effect sizes. Overall, the physical activity genes explained 16% (95% CI: 14.8, 19.8) of the variance in white blood cell count using a full model adjusted for fixed effects of age, sex, BMI, and smoking status and random effects of family and monozygosity. Physical activity genes significantly improved the goodness of fit, measured using a likelihood ratio test: χ^2^(30) = 192.84, p < 2.2 *e* ^−16^. Further, we rejected our null hypothesis of no enrichment of PA genes associated with WBC (hypergeometric P (x ≥ 13) = 0.000012).

We found one (1) PA gene, *SCN4A*, to be significantly associated (p < 0.05) with major depressive disorder, when the models were adjusted for age, sex, alcohol and the random effects of family and monozygosity. See Supplementary File A for detailed results.

### Association with Age Deceleration

We hypothesized that genes associated with physical activity are also associated with “healthy aging”, or, age deceleration (the deviation of chronological age from epigenetic or biological age) in diverse tissues, such as brain. We used 353 epigenetic markers^58^ to determine the epigenetic age of the individuals in the ROSMAP cohort in brain tissue, and verified the epigenetic age predictor was predictive of chronological age. The correlation between the estimated epigenetic age and chronological age was 0.69 with an absolute median deviance of 20 (See Supplementary File A). We took the difference between the epigenetic and chronological age to indicate age acceleration (“unhealthy aging”) or deceleration (“healthy aging”). We associated the difference between epigenetically predicted and chronological age with PA signature genes adjusting for sex and study (ROS vs MAP). Three PA signature genes, *PNPLA7, SSPN* and *NAPEPLD* were significantly associated (p < 0.05) and directionally concordant with age deceleration (see Figure 6 (a). Specifically, expression of *PNPLA7* and *SSPN*, up-regulated by physical activity, were associated with age deceleration, or equivalently, increased difference in chronological and biological age. A one SD unit increase in *SSPN*, Sarcospan, was associated with a 0.41-year (95% CI: 0.12, 0.71) deceleration in aging. On the other hand, increased expression *NAPEPLD*, down regulated by physical activity, was associated with a decrease in healthy aging - a one SD unit increase in *NAPEPLD* was associated with a 0.26 year decrease in healthy aging (95% CI: -0.45, -0.07). Overall, we found that the PA signature genes when adjusted for age and study explain 9% of the variance in age deceleration in the ROSMAP data with χ^2^(30) = 671.3, p < 0.042. The hypergeometric test was not significant (p = .12).

**Figure 6.**
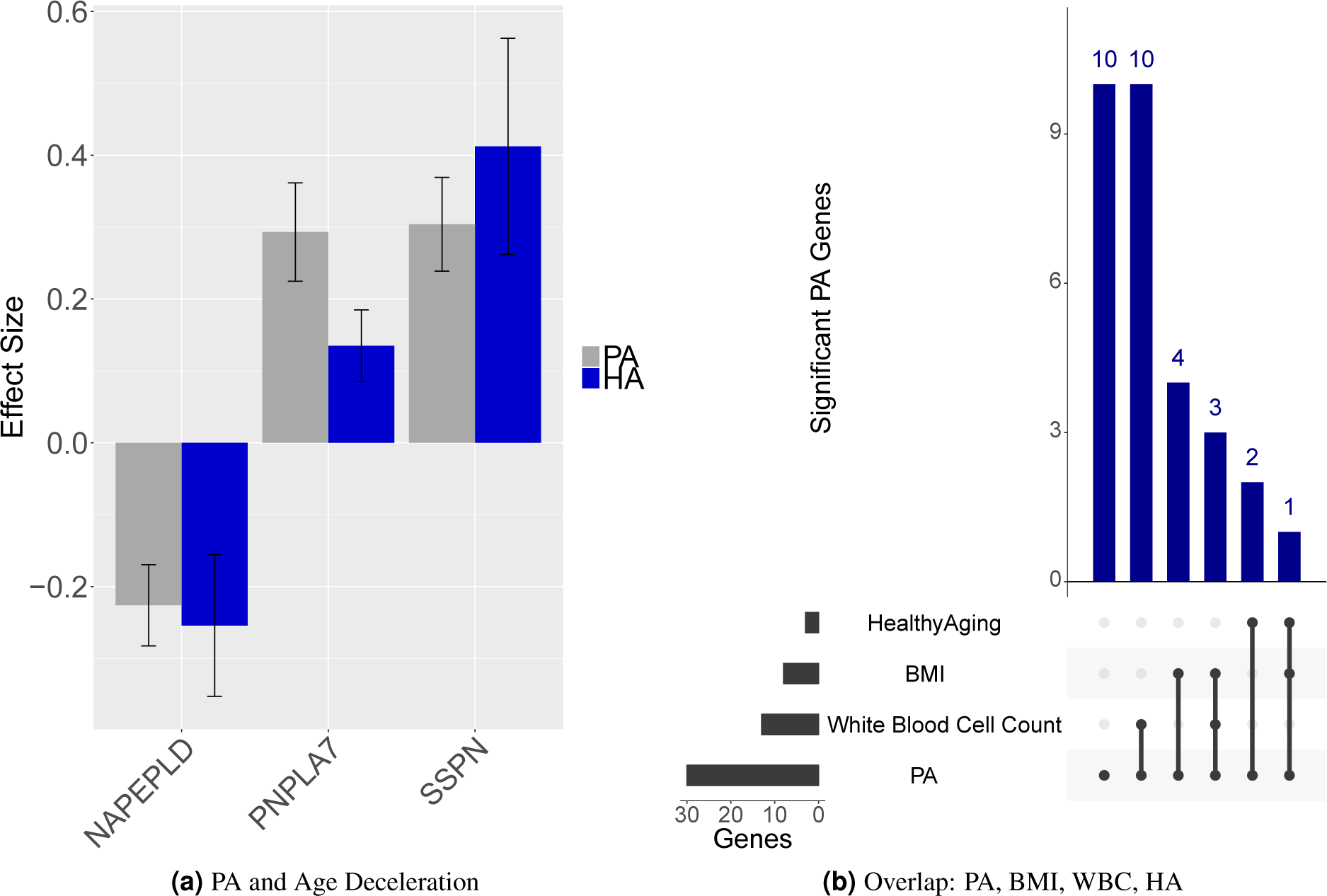
(a) Association of PA signature genes with age deceleration (healthy aging). (b) Overlap of PA signature genes across the three tested phenotypes, body mass index, white blood cell count and age deceleration.

Figure 6 (b) provides a summary of our key findings in terms of associations with the studied phenotypes: body mass index, white blood cell count and aging. We identified eight PA signature genes to be associated with BMI, thirteen associated with white blood cell count and three associated with aging. Of these YWHAZ(↓), ANXA1(↓) and CACGN1(↑) were associated with BMI and white blood cell count, while SSPN (↑) was associated with BMI in blood and aging in brain tissue.

## Discussion

Our work sought to identify changes in gene expression due to aerobic and anaerobic physical activity intervention by meta-analyzing 20 independent interventional gene expression studies. Second, we examined the biological relevance of the physical activity genes associated with differences in BMI, WBC, and MDD in monozygotic and dizygotic twins. Third, we examined the association between the PA signature and epigenetic aging of the brain. These 30 PA-associated genes have been implicated in different phenotypes in multiple GWASs and mouse model knockout systems. For example, *COL4A3*, part of the collagen trimerization and phospholipase-C pathways, has been linked to kidney diseases and cancers^71-72^, while *SLC4A5*, a sodium bicarbonate transporter has been associated with hypertension^73-75^. Overall, the PA genes have been associated with cardiovascular disease, adiposity, neurological, behavioral, and mortality and aging (See Supplementary File B for DisGeNET^76^ results). We describe specific associations in the following.

## Physical Activity associated gene expression and their correlation with BMI

We identified eight (8) PA signature genes, *C9orf131, CACNG1, EPS15L1, SSPN, ANXA1, SSX2IP, PNP* and *YWHAZ*, to be associated with body mass index in twins. All eight of these transcripts were directionally discordant with physical activity, that is transcripts up-regulated in physical activity were down-regulated in body mass index and vice versa. Transcripts *ANXA1, PNP, SSPN* and *YWHAZ* have all been previously associated with disorders of metabolism. *ANXA1*, Annexin A1, is a calcium and phospholipid-binding protein. It has been implicated in suppression of inflammation^77-80^ and is attenuated in obese individuals^81^. Akasheh^82^ and Warne^83^ et. al report ANXA1 deficient mice were leaner (as measured by epididymal fat pad weight) when compared to wild-type mice. Given our findings, we hypothesize that reduced activity of ANXA1 induced by physical activity may play a role in the reduction of adiposity.

Lim et al.^84,85^ have shown that YWHAZ depleted mice have improved glucose tolerance and significant reductions in adipose tissue, while overexpression of YWHAZ results in impaired insulin sensitivity and expansion of adipose tissue. We hypothesize that reduced YWHAZ expression induced by physical activity is associated with reduced adiposity.

High expression levels of PNP - a gene responsible for producing purine nucleoside phosphorylase enzyme - have been associated with higher purine content in the body^86^, a condition associated with increased risk of hyperuricemia and gout. Hyperuricemia and gout are known risk factors for cardiovascular disease, renal disease, insulin resistance, obesity and other metabolic diseases^87-90^. Given that decreased PNP is associated with decreased BMI in our twins cohort, we hypothesize that physical activity may play a role in the reduction of adiposity through regulation of the purine metabolism pathway. Heid et. al conducted a meta-analysis of 32 genome-wide association studies^91^ and identified SSPN as one of the 13 loci associated with waist-to-hip ratio and other measures of adiposity. More recently, Voisin et al. ^92^ and Keller et al.^93^ have identified SSPN, the sarcospan gene, to be significantly and negatively associated with BMI through epigenome-wide analysis. Based on these and our findings, we hypothesize that attenuation of SSPN induced by physical activity may reduce adiposity.

We found that aerobic and anaerobic physical activity alter the expression of a core set of genes. There is some evidence to support the role of both aerobic and anaerobic PA in decreasing risk for disease. For example, Sigal et. al^94^ in a randomized clinical trial to determine the optimal exercise modality for obese adolescents, found that body mass index (BMI) was decreased for aerobic groups (-1.0 (-1.7, -0.3)) and anaerobic groups (0.9(-1.7, -0.1)) but had a higher effect sizes for the combined exercise group (-2.0(-2.7, -1.2)). We claim that these two forms of exercise have similar routes for mitigation of disease risk, such as obesity.

## Physical Activity and White Blood Cell Count

We hypothesize that physical activity regulation of genes such as *ANXA1* may play a role in inflammatory-related phenotypes. For example, observationally, physical activity has been associated with reduced inflammation in the body^95,96^. In our investigation, we identified thirteen PA signature genes to be significantly associated and discordant in direction (12 out of 13 transcripts) with white blood cell count. Of these, *ANXA1, YWHAZ*, and *IL18R1* have been associated with inflammation in prior studies^79 97-103^. *ANXA1*, an anti-inflammatory protein annexin A1, for example, was found to be strongly expressed in Alzheimer’s disease and has been shown to promote resolution of inflammatory microglial activation^98^ and the clearance and degradation of amyloid-peptide^99^.

## Physical Activity and Age Deceleration in Brain Tissue

We identified three PA signature genes, *PNPLA7, SSPN*, and *NAPEPLD*, identified in blood/muscle tissue to be significantly associated (p < 0.05) and concordant in direction with age deceleration measured in brain tissue. These transcripts, *PNPLA7, SSPN*, and *NAPEPLD* are expressed in muscle, blood and brain tissue with *NAPEPLD* having high levels of expression in brain tissue^52^. *NAPEPLD*, N-acyl phosphatidylethanolamine phospholipase D, has been previously studied in the context of brain activity^104^ and *SSPN* has been associated with muscle regeneration^105^. We posit that physical activity regulation of *PNPLA7, SSPN*, and *NAPEPLD* in blood/muscle tissue is associated with age deceleration in the brain. This has significant clinical implications potentially providing a cost-effective and minimally invasive mechanism for measuring age deceleration.

## Limitations

While, to our knowledge, we have reported largest study to-date in gene expression differences in physical activity, our study has several limitations. First, our meta-analysis spans a broad range of age groups, from 8 year olds to individuals in their 60s. While we have identified transcripts that are significantly differentially expressed across this age span, we may have potentially missed some age specific signatures of physical activity. In a similar vein, our meta-analysis does not focus on sex-specific regulation of genes induced by physical activity. Second, while we ensured uniformity of physical activity intervention with respect to time duration, we could not fully ensure uniform intervention (for example, type of strengthening exercises, equipment used etc.). This could potentially alter effect sizes/significance of identified genes. Third, we used both blood and muscle tissues in our meta-analysis (2/3 of the tissue was blood). This could potentially bias our results perhaps warranting additional analysis with more muscle samples, and other pathologically relevant tissues, such as liver, adipose, and brain. Fourth, our population for the meta-analysis is largely homogeneous (non-Hispanic white, based on the studies that reported ethnicity). Our discovered PA associated genes may not hold true for a heterogeneous population. These results must be validated in an ethnically diverse population to be generally applicable. Fifth, we make extrapolations in aging associations in brain tissue, when our expression signature was derived from blood tissue. Others have documented modest to null correlation of blood gene expression to that in brain, so our brain-related associations require replication. Sixth, we found little association between major depressive disorder and physical activity. This could potentially be due to the small number of cases in our twin cohort and requires replication in a larger cohort.

## Conclusions

Our analysis show that physical activity induces differentiation of thirty genes, with 20 transcripts over-expressed and 10 transcripts down-regulated. Further, we find several of the PA significant genes are associated with body mass index, white blood cell count, and age deceleration. We find that phenotype-associated genes up-regulated by physical activity were found to be down-regulated in body mass index and white blood cell count, and up-regulated in age deceleration and vice versa. These findings establish a directionally consistent molecular link between physical activity and these phenotypes, under-scoring the potential therapeutic benefits of physical activity.

There are several outstanding questions. While our results highlight the common molecular effects of aerobic and anaerobic exercise, the differences in the molecular signature induced by aerobic vs anaerobic exercise and their biomedical relevance is an open question. There have been prior observational studies detailing the effect of exercise modality on phenotypes (waist circumference, percentage of body fat)^94 106^ but the changes in the gene expression caused by the different exercise modalities and their association with body mass index or aging has not been studied. Second, while our work has measured the genes modulated via physical activity, it is unclear how long, in terms of time duration, these modifications to the gene expressions persist. Stratifying gene expressions by the minutes/hours of physical activity per week/month can provide potential insights into this question and potentially a molecular basis for recommendations on daily/weekly levels of physical activity. Third, while we have identified PA signature genes, additional validation in an independent cohort is required to develop a generalized PA predictor to determine the physical activity in individuals. Fourth, physical activity is a therapeutic for many chronic conditions. We have examined the associations between physical activity and body mass index, white blood cell count and healthy aging. Further investigations are needed to identify molecular level associations of physical activity with other chronic conditions such as Alzheimer’s disease.

